# The impact of modern jazz dance on the electrical brain activity

**DOI:** 10.1101/472324

**Authors:** Johanna Wind, Wolfgang Schöllhorn

## Abstract

Dance as one of the earliest cultural assets of mankind is practised in different cultures, mostly for wellbeing or for treating psycho-physiological disorders like Parkinson, depression, autism. However, the underlying neurophysiological mechanisms are still unclear and only few studies address the effects of particular dance styles. For a first impression, we were interested in the effects of modern jazz dance (MJD) on the brain activation that would contribute to the understanding of these mechanisms. 11 female subjects rehearsed a MJD choreography for three weeks (1h per week) and passed electroencephalographic (EEG) measurements in a crossover-design thereafter. The objectives were to establish the differences between dancing physically and participating just mentally with or without music. Therefore, each subject realized the four following test conditions: dancing physically to and without music, dancing mentally to and without music. Each of the conditions were performed for 15 minutes. Before and after each condition, the EEG activities were recorded under resting conditions (2 min. eyes-open, 2 min. eyes-closed) followed by a subsequent wash-out phase of 10 minutes.

The results of the study revealed no time effects for the mental dancing conditions, either to or without music. An increased electrical brain activation was followed by the physical dancing conditions with and without music for the theta, alpha-1, alpha-2, beta and gamma frequency band across the entire scalp. Especially the higher frequencies (alpha-2, beta, gamma) showed increased brain activation across all brain areas. Higher brain activities for the physical dancing conditions were identified in comparison to the mental dancing condition. No statistically significant differences could be found as to dancing to or without music. Our findings demonstrate evidence for the immediate influence of modern jazz dance and its sweeping effects on all brain areas for all measured frequency bands, when dancing physically. In comparison, dancing just mentally does not result in similar effects.

## Introduction

The human ability to dance can be traced back as far as human bipedal walking, in other words, about 2–5 million years ago. Cave paintings indicate whole body movements as used in dance [1–3]. Independent of the fascination emanating from dance in all cultures, dance is differentiated into numerous categories, such as classical dance, modern dance, folk dance, spiritual dance, etc. The amalgamation of jazz and modern dance does not only lead to a new, particular dance style, called modern jazz dance, that receives increasing attractiveness in the European dance community [4].

The specific techniques of jazz dance are mainly characterized by powerful and dynamic movements with elements of isolation, poly-centricity and poly-rhythmicity [5]. The modern dance technique is typically associated with the principles of >contraction and release< as well as >fall and recovery< [6]. Most frequently applied metaphors are also >rebound<, >swing<, >suspension< and >off balance< [7] in the context of modern dance.

Besides the pleasure of watching dancers at theatre performances or dancing contests, dance receives a growing interest for health reasons. Some studies indicate improvements when dance is used alongside medical therapies for breast cancer [8, 9], diabetes [10], fall prevention [11, 12] or dementia [13, 14]. An augmentation of the effects could be established when implementing dance therapy in the treatments for autism [15], depression [16–18] and Parkinson [19–22]. In its multifaceted ways, dance is assumed to promote human creativity [23] or can contribute to raising wellbeing [24] and can lead to the experience of flow [25].

More recently, the underlying neurophysiological mechanisms of dancing moved into the focus of interest. The effect of dance observation and its influence on EEG brain activity was studied and the findings recorded an increased activation of the premotor cortex [26, 27]. Cross, Hamilton and Crafton [28] investigated professional dancers, with functional magnetic resonance imaging (fMRI) to fathom the brain activity in more detail during the action of dancing. The dancers underwent whole-body dance training for five weeks (5 h a week), whereby EEG-recordings were taken at the end of each week. During the fMRI-measurements, the dancers observed the rehearsed and unrehearsed movements of a model dancer. In comparison to dance movements that were unrehearsed, the premotor and parietal areas were enhanced during the observation of rehearsed movements which the dancers were able to execute.

Another fMRI study [29] showed differences of the gender brain activity in professional dancers. It exists in gender specific movements observed by one gender during the training sessions without executing them. Increased brain activity for the premotor, parietal and cerebellar could be revealed when female and male dancers watched gender-specific movements. These effects were not observed when watching gender-different dancers. Poikonen, Toiviainen and Tervaniemi [30] compared the brain activity by means of EEG analysis gained from dancers, musicians and laymen. They detected theta-synchronisation at the fronto-central electrodes when dancers watched an audio-visual sequence of the choreography for *Carmen.* This effect was neither identified in the musicians nor the laymen. During fast movements, the alpha frequency decreased for all groups assumingly because of the increased mental effort. An event-related desynchronization in the alpha and lower beta frequencies was observed by Orgs, Dombrowski and Jansen-Osmann [31] when professional dancers watched dance movements in contrast to everyday movements. Dancers showed a reduction in the alpha and beta band frequency when watching dance movements, however, non-dancers showed no decrease. Fink, Graif and Neubauer [32] focused on the EEG effects of a mentally self-developed (improvised) dance in their study, more specifically on the recall of a familiar dance style. They compared the EEG brain activity of professional dancers and dance novices while imagining a very creative, improvised dance, on the one hand, and a traditional waltz, on the other hand. No differences were found between the two groups for the waltz task, though for the improvisation task, the professional dancers showed increased right-hemispheric alpha frequencies. Thus, there is evidence for coherence between the right-hemispheric alpha synchronization and creativity.

Indeed, these studies showed support for various effects of dance observations. The influence of physically self-executed, during and immediately after, dance movements is a rarer research objective.

Commonalities and differences between the mental and active learning of a dance was investigated by Cross et al. [33]. They measured common active brain areas with fMRI just as Cross, Hamilton and Grafton had done [28] during the process of learning a techno dance sequence through observation only and also by physically learning it. After five days of rehearsal, the premotor and parietal brain areas were more activated than before. A study by Müller et al. [34] investigated two groups of elderly non-dancers (68-80 years) over a period of six to 18 months. The first group realized dance training sessions and the second group a fitness training. After six months, the MRI-test provided evidence for increased grey matter volume at the gyrus praecentralis and an increase at the gyrus parahippocampalis, after 18 month for the dance training group only.

Similar to the study by Fink et al. [32], an increased alpha activity and beta frequency could be observed for professional dancers in the study by Ermutlu et al. [35]. They compared dancers’ with fastball athletes’ brain activity as well as that of a control group. The measurements referred to the resting condition.

According to current knowledge, the PET study by Brown, Martinez and Parsons [36] is one of the few studies which measured the brain activity while dancing. They attended a step sequence of a tango and managed to figure out the activation of the anterior cerebellum vermis lasting the entrainment of tango steps to musical accompaniment. Metric Tango steps to a regular, metric rhythm led to an increased activation at the right putamen in comparison to an irregular rhythm.

Cruz-Garza et al. [37] examined five dancers experienced in Laban movement notation with EEG and inertial sensors while performing three different manners: moving in a non-intentional way, thinking about an intentional movement and dancing the imagined intentional movement. Just thinking about an expressive movement activated the prefrontal, motor, and parietal areas. At least, when dancing this expressive-thought dance, all aforementioned brain areas increased in activation.

In sum, the premotor cortex, parietal areas and the cerebellar revealed more activation while observing dance movements in comparison to the baseline brain activity [26–29]. Moreover, an increased right-hemispheric alpha frequency could be noted at the parietotemporal and parietooccipital areas for professional dancers and even in regards to the beta band [32, 35]. Contrastingly, Orgs et al. [31] could show a decrease of the alpha- and beta band frequencies in professional dancers while watching dance movements. In the case of acquiring the dance sequences physically, before or while actually dancing, the prefrontal, motor, parietal areas, the anterior cerebellum vermis and the right putamen increased in activation [36, 37]. Furthermore, the grey matter gained in volume at the gyrus praecentralis and the gyrus parahippocampalis [34].

However, only few studies examined whole-body movements and determined a specific dance style. The actual or immediate effects of physical dance were investigated in their entirely in the studies by Brown et al. [36] in a tango sequence and Cruz-Garza et al. [37] in dancing expressive movements. No studies about the effects of modern Jazz Dance and the different reactions as to physical and mental dance or the specific effect of music inclusion or non-inclusion could be found.

The aim of the current study is to examine the different effects on the spontaneous electrical brain activity caused by physical dance or the imagination of dance, with and without music.

## Material and Methods

### Participants

Eleven female subjects with a mean age of 24.3 years (*SD* = 2.45; range: 21-29) volunteered for this study. The subjects were recruited from the Johannes Gutenberg University of Mainz and did not have to meet any eligibility requirements. The inclusion criteria for the study was finishing three modern jazz dance training sessions of 1 hour each and having no experience in any dance style. All subjects were healthy, free from neurological diseases and right-handed. Five of eleven subjects ingested the birth control pill daily. The constraints as to only female subjects is ascribed to the gender-specific differences of the brain [38, 39]. The elucidation as regards the purpose of the study and the informed consent from the subjects was given. The local ethics committee of the Johannes Gutenberg University of Mainz (Germany) approved the study.

### Experimental Procedure

The electroencephalography (EEG) measurements took place in a dimly lit room, in which all training conditions were conducted. Each testing condition was preceded and followed by a resting condition (2 min. eyes-open, 2 min. eyes-closed) with a subsequent wash-out phase of 10 minutes. During the resting and wash-out phases the subjects were asked to sit calmly on a chair facing a white wall.

All subjects accomplished each testing condition for the modern jazz dance (MJD) choreography in a within-subject design. Four testing conditions had to be passed in a random sequence: dance the MJD-choreography physically with music (pwm), physically with no music (pnm), mentally with music (mwm) as well as mentally with no music (mnm).

Three weeks before the EEG measurements were taken, all subjects attended a once a week training course in modern jazz dance (in total three times) in the sports facility of the Johannes Gutenberg University Mainz. All training courses were the same for all subjects. Three to five days after the last training session, the EEG measurements were taken. All testing conditions were executed with eyes open, even the mental ones.

### EEG Data Acquisition and Analysis

The EEG measurements were recorded with the Micromed SD LTM 32 BS amplifier (Venice, Italy) and the System Evolution Plus Software (Venice, Italy). Nineteen electrodes (Fp1, Fp2, F3, F7, Fz, F4, F8, C3, Cz, C4, T3, T4, P3, P7, Pz, P4, P8, O1, O2) were applied according to the international 10-20-system with the reference electrode attached to the nose. The electrode impedances were kept below 10 kΩ. The EEG signals were digitized at a sampling rate of 1024 Hz with a bandpass filter from 0.008 Hz to 120 Hz. The electrooculography (EOG) was affixed at the lateral orbital and the medial upper rim of the right eye. The spontaneous EEG and EOG recordings (2 min. eyes-open condition, 2 min. eyes-closed) were inspected to different artefacts (e.g. muscle contractions), which were removed in the end. To avoid the increase of alpha waves due to the closing of the eyes, we merely analyzed the 2 minutes eyes-open condition [40]. With the aid of a Fast-Fourier analysis, the mean power amplitudes were obtained in theta (3.5-7.5 Hz), alpha-1 (7.5-10 Hz), alpha-2 (10-12.5 Hz), beta (12.5-30 Hz) and gamma (30-70 Hz). These frequency ranges were progressed by Zschocke and Hansen [41].

### Statistical Analysis

The within-subject factors were analyzed by means of a three-factor repeated-measure analysis of variance (ANOVA) and Bonferroni corrected post hoc tests. Factor one comprised the four testing conditions MJD pwm, pnm, mwm, and mnm, repeated measures were the second factor with two time conditions (pre-test, post-test) and the electrode positions constituted the third factor (Fp1, Fp2, F3, F7, Fz, F4, F8, C3, Cz, C4, T3, T4, P3, P7, Pz, P4, P8, O1, O2). The statistical significance was set at p-value ≤ 0.05. Additionally, the effect size (Cohen’s *η*^2^, 1988) was particularized, with the following conventions: *η*^2^ = 0.01 (small effect), *η*^2^ = 0.06 (medium effect), *η*^2^ = 0.14 (large effect).

## Results

### Statistical Analysis EEG

Fig 1 shows the mean power of the theta, alpha-1, alpha-2, beta and gamma frequencies after the four specific testing conditions. The frequency scales are accommodated to the corresponding frequency bands. Table 1 presents the significant p-values for electrode positions of physical dancing to (pwm) and without (pnm) music at the post-tests and furthermore for statistical significant differences between the testing conditions.

**Fig 1.**
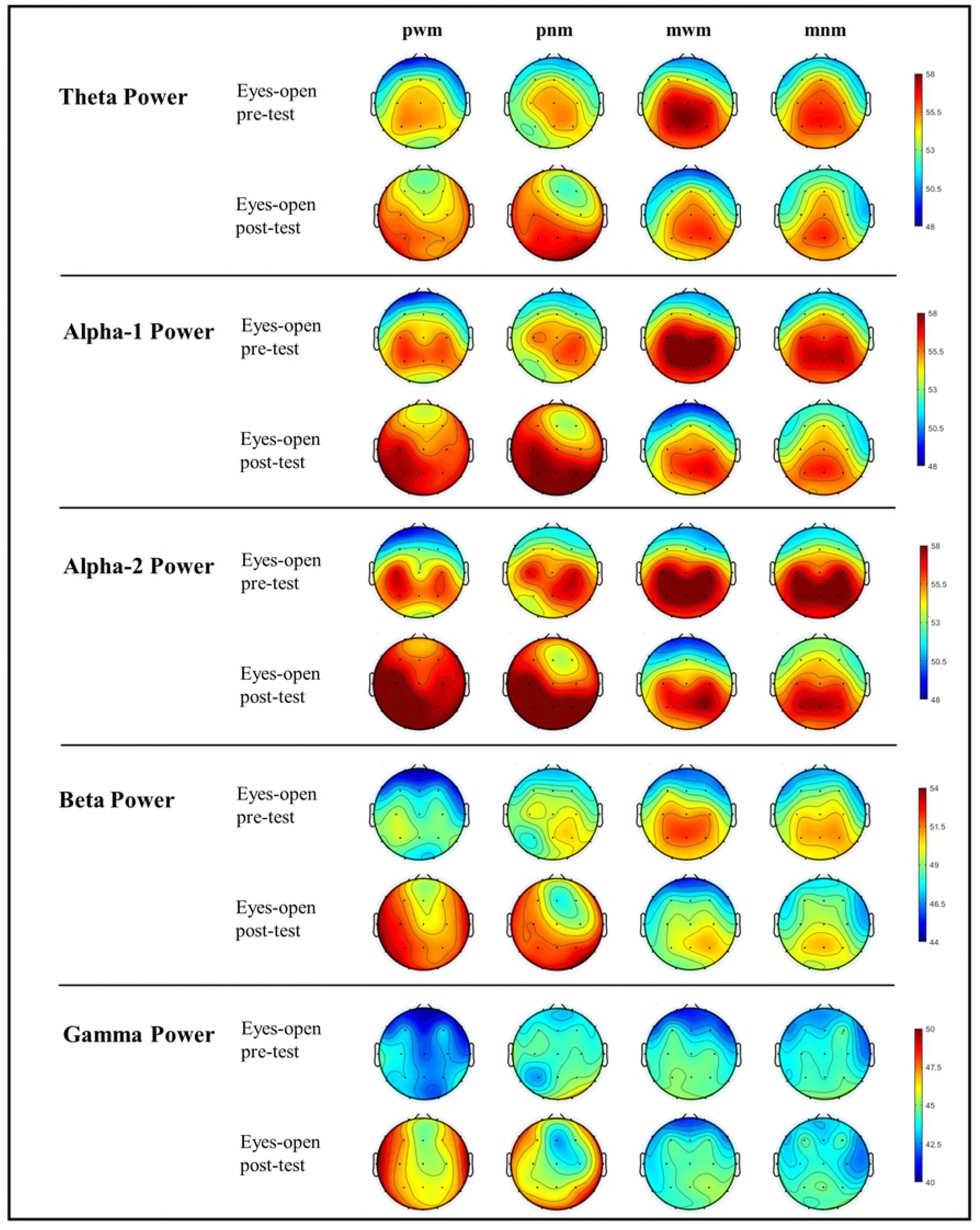
Spontaneous brain activity across all frequencies, four testing conditions, pre- and post-test. All frequency bands showed an increase after physical dancing to and without music for the eyes-open condition. No significant time effects were found for mental dancing with and without music.

**Table 1.** Significant p-values for the electrode positions of all testing conditions. Only the testing conditions physical dancing to (pwm) and without (pnm) music revealed significant p-values at the post-test. All frequency bands showed significant p-values at the comparison of the four testing conditions.

| power spectrum | testing conditions (in comparison) | electrode position with p-value |  |  |  |  |
| --- | --- | --- | --- | --- | --- | --- |
| theta | pwm | Fp1 ($p = 0.022$) | F8 ($p = 0.015$) | T3 ($p = 0.010$), | T4 ($p = 0.009$) | O1 ($p = 0.048$) |
| | pnm | Fp2 ($p = 0.044$) | O1 ($p = 0.025$) | O2 ($p = 0.017$) | | |
| | (pwm/mwm)<sup>a</sup> | T3 ($p = 0.028$) | T4 ($p = 0.002$) | | | |
| | (pnm/mnm)<sup>d</sup> | T4 ($p = 0.015$) | | | | |
| alpha-1 | pwm | Fp1 ($p = 0.008$)<br>O1 ($p = 0.012$) | F7 ($p = 0.009$)<br>O2 ($p = 0.016$) | F8 ($p = 0.026$) | T3 ($p = 0.009$) | T4 ($p = 0.019$) |
| | pnm | Fp1 ($p = 0.032$) | Fp2 ($p = 0.024$) | F7 ($p = 0.032$) | O1 ($p = 0.016$) | O2 ($p = 0.009$) |
| | (pwm/mwm)<sup>a</sup> | F8 ($p = 0.046$) | T3 ($p = 0.024$) | T4 ($p = 0.022$) | | |
| | (pwm/mnm)<sup>b</sup> | F8 ($p = 0.045$) | | | | |
| | (pnm/mwm)<sup>c</sup> | F7 ($p = 0.035$) | O2 ($p = 0.040$) | | | |
| | (pnm/mnm)<sup>d</sup> | T4 ($p = 0.006$) | O2 ($p = 0.026$) | | | |
| alpha-2 | pwm | Fp1 ($p = 0.003$)<br>F4 ($p = 0.020$)<br>Pz ($p = 0.043$) | Fp2 ($p = 0.023$)<br>F8 ($p = 0.010$)<br>T6 ($p = 0.025$) | F7 ($p = 0.001$)<br>T3 ($p = 0.003$)<br>O1 ($p = 0.001$) | F3 ($p = 0.28$)<br>T4 ($p = 0.013$)<br>O2 ($p = 0.002$) | Fz ($p = 0.044$)<br>T5 ($p = 0.007$) |
| | pnm | Fp1 ($p = 0.045$)<br>O1 ($p = 0.016$) | Fp2 ($p = 0.035$)<br>O2 ($p = 0.007$) | F7 ($p = 0.042$) | F8 ($p = 0.047$) | T6 ($p = 0.038$) |
| | (pwm/mwm)<sup>a</sup> | Fp1 ($p = 0.028$)<br>O1 ($p = 0.004$) | F7 ($p = 0.027$) | F8 ($p = 0.024$) | T3 ($p = 0.004$) | T4 ($p = 0.048$) |
| | (pwm/mnm)<sup>b</sup> | F7 ($p = 0.031$) | F8 ($p = 0.034$) | O1 ($p = 0.046$) | | |
| | (pnm/mwm)<sup>c</sup> | O2 ($p = 0.047$) | | | | |
| | (pnm/mnm)<sup>d</sup> | T4 ($p = 0.039$) | O2 ($p = 0.021$) | | | |
| beta | pwm | Fp1 ($p = 0.008$)<br>T3 ($p = 0.005$)<br>T6 ($p = 0.012$) | Fp2 ($p = 0.048$)<br>C3 ($p = 0.018$)<br>O1 ($p = 0.009$) | F7 ($p = 0.001$)<br>T4 ($p = 0.001$)<br>O2 ($p = 0.004$) | F3 ($p = 0.046$)<br>T5 ($p = 0.008$) | F8 ($p = 0.004$)<br>Pz ($p = 0.048$) |
| | pnm | O2 ($p = 0.030$) | | | | |
| | (pwm/mwm)<sup>a</sup> | F7 ($p = 0.021$) | T3 ($p = 0.014$) | T4 ($p = 0.014$) | | |
| | (pwm/mnm)<sup>b</sup> | F8 ($p = 0.023$) | T4 ($p = 0.027$) | | | |
| | (pnm/mnm)<sup>d</sup> | T4 ($p = 0.007$) | O2 ($p = 0.025$) | | | |
| gamma | pwm | Fp1 ($p = 0.019$)<br>F8 ($p = 0.001$)<br>T4 ($p = 0.001$)<br>T6 ($p = 0.029$) | Fp2 ($p = 0.047$)<br>T3 ($p = 0.005$)<br>T5 ($p = 0.006$)<br>O1 ($p = 0.040$) | F7 ($p = 0.001$)<br>C3 ($p = 0.014$)<br>Pz ($p = 0.021$)<br>O2 ($p = 0.008$) | F3 ($p = 0.030$)<br>Cz ($p = 0.012$)<br>P3 ($p = 0.012$) | Fz ($p = 0.035$)<br>C4 ($p = 0.045$)<br>P4 ($p = 0.011$) |

<sup>a</sup> comparison between the testing conditions physical with music (pwm) and mentally with music (mwm)
<sup>b</sup> comparison between the testing conditions physical with music (pwm) and mentally non music (mnm)
<sup>c</sup> comparison between the testing conditions physical non music (pnm) and mentally with music (mwm)
<sup>d</sup> comparison between the testing conditions physical non music (pnm) and mentally non music (mnm)

The ANOVA of the theta frequency data revealed a significant interaction effect for testing x time x electrodes (*F*(54,540) = 1.473, *p* = 0.019, η^2^ = 0.128). Post hoc comparisons showed significant differences at the post-test between pwm and mwm for the electrode positions T3 and T4, whereby pwm was accompanied by higher electrical brain activity. Significant differences between pnm and mnm for the electrode position T4 could be found. Only the testing conditions pwm showed a time effect with an increased mean power for the electrode positions Fp1, F8, T3, T4, O1 and the testing condition pnm for the electrodes Fp2, O1, O2 (for p-values, see Table 1).

The ANOVA of the alpha-1 frequency band data showed an interaction effect for testing x time, (*F*(3,30) = 4.096, *p* = 0.015, η^2^= 0.291) and an interaction effect for testing x time x electrodes, (*F*(54,540) = 1.461, *p* = 0.021, η^2^= 0.128). Post hoc comparisons revealed significantly increased power after the testing condition pwm compared to mwm for electrodes F8, T3, T4 and compared to mnm for the electrode position F8.

Furthermore, increased power after pnm testing compared to mwm testing was identified for the electrode positions F7 and O2. The pnm testing also showed increased power for electrodes T4 and O2 in comparison with the mnm testing condition. The testing conditions pwm led to increased power following the testing for the electrode positions Fp1, F7, F8, T3, T4, O1, O2 and the pnm testing for the electrode positions Fp1, Fp2, F7, O1, O2 (for p-values, see Table 1).

The ANOVA of the alpha-2 power revealed a significant effect of time, (*F*(1,10) = 7263, *p* = 0.023, η^2^ = 0.421), and an interaction effect between testing x time, (*F*(3,30) = 5,176, *p* = 0.005, η^2^ = 0.341). An interaction effect between testing x time x electrode positions, (*F*(54,540) = 3.401, *p* = 0.010, η^2^ = 0.134) was also detected.

Post hoc comparisons showed an increased alpha-2 power after the training condition pwm compared to the mwm testing condition for electrodes Fp1, F7, F8, T3, T4, O1 and to the mnm testing for electrodes F7, F8, O1. The pnm testing conditions presented increased power at the post-test for the electrode positions T4 and O2 in comparison to the mnm testing and for electrode O2 at the mwm testing (for p-values, see Table 1). Likewise, for the alpha-1 frequency, alpha-2 frequency revealed a time effect only for the testing conditions pwm and pnm. The post hoc test indicated increased power for electrodes Fp1, Fp2, F7, F3, Fz, F4, F8, T3, T4, T5, Pz, T6, O1 and O2 after the pwm testing condition. Also, the pnm testing conditions showed increases of the power for the electrode positions Fp1, Fp2, F7, F8, T6, O1 and O2 (for p-values, see Table 1).

The ANOVA for the beta-frequency band led to significant differences between the pre- and post-tests, (*F*(1,10) = 8,457, *p* = 0.016, η^2^ = 0.458). There was also a significant interaction when testing x time, (*F*(3,30) = 3.836, *p* = 0.019, η^2^ = 0.277). No interaction effect for testing x time x electrodes was found, however, post hoc comparisons showed significant differences.

The analysis also revealed an increased brain activation after the pwm condition compared to the mwm condition for the electrode positions F7, T3, T4 and compared to the mnm condition for electrodes F8, T4.

The pnm condition effected increased power for electrodes T4 and O2 in comparison to the mnm testing. An increased effect was also noticeable from pre- to the post-test for the pwm condition for several electrode positions, Fp1, Fp2, F7, F3, F8, T3, C3, T4, T5, T6, Pz, O1, O2. The pnm condition revealed increased power after the testing for the electrode position O2 (for p-values, see Table 1).

The ANOVA for the gamma band revealed an effect of time, *F*(1,10) = 12.088, *p* = 0.006, η^2^ = 0.547. Again, there was no significant interaction effect of testing x time x electrodes. However, post hoc comparisons showed increased power after the pwm condition for the electrode positions, Fp1, Fp2, F7, F3, Fz, F8, T3, C3, Cz, C4, T4, T5, T6, Pz, P3, P4, O1 and O2 (for p-values, see Table 1).

### Discussion

The aim of this study was to investigate the effects of dancing a modern jazz choreography in four different modes on the electrical brain activity. The four modes were dancing the choreography physically or mentally, with or without music. The subjects were inexperienced dancers and had to learn a modern jazz dance choreography within three weeks (1h per week sessions) before the actual EEG measurements were recorded. Every subject passed the testing conditions (pwm, pnm, mwm, mnm) consecutively on the same day with a 15-minute wash-out phase between each condition. The EEG’s were measured immediately before and after the particular training condition, in terms of a resting condition (2 minutes eyes-open, 2 minutes eyes-closed). To avoid the increase of alpha waves due to the closing of the eyes, we focused the analysis on the 2 minutes eyes-open condition [40].

In the present study, we could assess an increased power across all frequency bands when the subjects danced physically to or without music in comparison to the mental dance conditions. Especially, the condition “physical dance with music” (pwm) led to an increased power of the alpha-2, beta and gamma frequencies at the frontal, temporal and occipital brain areas during post-test. Compared to the theta, alpha-1, alpha-2 and beta frequency bands, the gamma frequency showed increased power at the central and parietal brain areas. No time effects were observed for the mental testing conditions mwm and mnm.

These results are in accordance with the findings of the studies previously mentioned. Similar to Fink et al. [32] and Ermutlu et al. [35], we could show a higher activation of the alpha and beta bands in comparison to the baseline after the physical dance. In difference to Fink et al. [32] and Ermutlu et al. [35], the present study could reveal the aforesaid higher activations for unexperienced dance subjects, not for the mentally (improvised) dancing [32], but rather for the physical dancing. Fink et al. [32] could show an increased right-hemispheric alpha frequency for professional dancers when mentally dancing an improvisation dance. Ermutlu et al. [35] could reveal an increased alpha and furthermore a beta band activation for professional dancers as could Fink et al. [32]. As regards the physical dance conditions, we furthermore detected increased gamma power across all brain lobes and theta power for electrodes Fp1, Fp2, F8, T3, T4, O1, O2. Whether this increase is due to the chosen choreography or the level of the dancers demands for further research.

Merely, Poikonen et al. [30] revealed a decrease of the alpha phase synchrony for several electrode pairs across the brain for dancers while watching an audio-visual choreography of Bizet’s *Carmen.* They linked the decrease to increased attention because of the fast movements. In addition, Orgs et al. [31] demonstrated a decrease for the alpha and lower beta frequency bands for professional dancers while watching dance movements in comparison to everyday movements.

The fMRI-studies by Cross et al. [28], Cross et al. [33] and the EEG-study by Cruz-Garza et al. [37] showed an increased activation of the parietal brain areas. For the study by Cruz-Garza et al. [37], the parietal areas were activated at the delta frequency. In comparison to the present study, Cross et al. [28] revealed the higher activation of the parietal areas while observing rehearsed dance movements in an fMRI study. Cross et al. [33] presented the increase not only for learning a techno dance sequence through observation, but also by learning it physically. Next to the higher activation of the parietal areas, Cruz-Garza et al. [37] revealed a higher activation of the prefrontal cortex while performing an expressive dance as well. The current study indicated a higher activation of the parietal brain areas, too, but compared to Cross et al. [28, 33], this was detected immediately after physically dancing a modern jazz dance to and without music for the alpha-2, beta and gamma frequency bands. Like Cruz-Garza et al. [37], the current study showed increased power at the prefrontal cortex immediately after the dancing for all frequency bands.

The findings of the present study and those ascertained by Cross et al. [28], Cross et al. [33] and Cruz-Garza et al. [37] can be taken as indicators for the activation of the same brain areas of dance observation, the immediate effects following the dancing, and while dancing.

However, there is evidence for a significant difference between dancing physically and imagining a previously learned choreography, at least in beginners. We may not have checked objectively whether the subjects imagined the actual required choreography, so it is possible that they were thinking about different things, which could have obstructed any higher brain activation. Nevertheless, it also might have been the case that just imaging a dance choreography leads to no significant changes in the electrical activity. The study by Ott [42] provides evidence for increased brain activation, predominantly in alpha and theta frequency bands, after physical yoga in comparison to only meditative yoga. These results substantiate the findings of this study, in the manner that physical activity more significantly leads to alterations in the brain. For this reason, we do not discuss the mental dancing condition in more detail, but rather parse the physical dancing conditions.

For the pwm and pnm conditions, the theta and the alpha-1 frequencies were activated at fewer electrode positions, compared to the higher frequency bands. Theta activity is primarily associated with a drowsy state and appears more often in childhood than adulthood [43]. Furthermore, Malik and Amin [43] found an increased theta activity in attentional processing and working memory as well. Especially frontal midline theta is associated with working memory, processes of anxiety and cognitive control [44, 45]. Indeed, the current study presented a time effect for conditions pwm and pnm, but solely for the electrode positions Fp1, Fp2, F8, T3, T4, O1 and O2. The activation of the frontal lobe is also connected to the Brodmann areas (BA) 10, 46, 9 and 45, which signify i.a. to working memory [46]. Thus, the increased activation of the theta frequency at the prefrontal cortex could be a cue for the involvement of the working memory from cognitive psychology. Awareness processes are also linked to the frontal lobe [47] and could happen in connection with attentional processes lasting an increased theta activity. Whether the increased theta frequencies after physical dancing influence healing processes [15–22] in the form of increased dopamine production [48] or by activating the parasympathetic system [49] or supporting the absorption of nutrients [50] needs to be investigated in detail. Theory already provides plausible evidence.

Even for the alpha-1 frequency band, few electrode positions are activated at the frontopolar, frontal temporal and occipital brain areas after physically dancing to and without music (see for details Table 1). The alpha state (7,5-12,5 Hz) is often related to psychic and physical relaxation while still maintaining vigilance [41, 43]. Alpha increases especially when no mental task is to be performed and occurs more often in the occipital and parietal areas [41, 43]. Cantero, Atienza and Salas [51] proved an increase of slower alpha waves in the anterior brain areas and an activation peak in the occipital area. Accordingly, the fronto-central alpha pattern is associated with a typical feature of drowsiness. As per Klimesch [52] and Hanslmayr et al. [53], the alpha frequency is assumed to be the basis of memory and attention processes, which are processed in the frontal lobe. These frequency and area specific properties could be an indication for an attentional process while dancing to and without music because of the recall of the previously rehearsed modern jazz dance choreography. In addition to this, it has been suggested that the attention in combination with the activated temporal lobe incorporates the reprocessing and perception of the heard music [54, 55]. It could be furthermore an indication for reprocessing the triggered emotions due to the music. The occipital brain activity mirrors the vision, on the one hand, and on the other hand, it may be evidence for proprioceptive attention while physically dancing [55–57]. The subjects had to concentrate on their limbs the whole time, to coordinate them in the required manner. This procedure needs proprioceptive attention. The assertions proclaimed by Zschocke and Hansen [41] and Cantero et al. [51] about alpha in sum and particularly slower alpha waves fit the present findings. The conditions pwm and pnm showed higher activations in the posterior brain regions and additionally the anterior. As per Cantero et al. [51], the simultaneous appearance of anterior and posterior brain activation denotes a relaxed wakefulness, which might be evidence for the relaxed and wakeful states of the subjects after the physical dancing condition. Henz’s and Schöllhorn’s study [58] investigated the impact of the physical and mental Qigong technique Wu Qin Xi. They revealed i.a. a shift in alpha-1 and alpha-2 from posterior to anterior brain regions after physically exercising the Qigong technique. Similar activation patterns of the brain were already found after 10 minutes of differential training, where a gross motor movement technique was trained without repetition and without augmented feedback [59]. Both studies go along with and extend the “transient hypofrontality hypothesis”, which suggests that moderate, aerobic range, exercises a result in a concomitant transient decrease of the activity of the prefrontal cortex. [60]. In addition to long-lasting, cyclic endurance sports, meditation, hypnosis or dreaming are also supposed to cause similar decreased brain activations in the frontal area [60, 61]. This altered brain activation is often associated with the change of consciousness and can lead to a trance-like state (diminished awareness of surroundings, timelessness, living in the here and now, peacefulness, floating). Our results suggest that physical dances of shorter duration compared to something typically seen in endurance sports, with and without music, can lead to a down regulation of the frontal lobe. The increased alpha band frequencies at the prefrontal electrodes show similar features to those explained in the transient hypofrontality hypothesis. Assumed the subjects dance the choreography more automatically, the more the alpha activity is increased at the prefrontal cortex and further afield, the more they reach a higher state of consciousness. Whether these higher states of consciousness due to dance are one reason for the attractiveness of folk and group dances and their survival in different cultures over thousands of years, and whether the effects are amplified in groups, is still speculation at this moment and needs further research. But the present findings in addition to existing literature provides strong evidence for this cultural phenomenon.

Higher alpha frequencies are also linked to creative thinking when imagining an improvisation dance, if the prefrontal areas are activated [23]. Similar to Fink et al. [23], our study revealed high alpha waves at the prefrontal brain areas, too, however not while imagining an improvised dance, but rather after actual dancing a modern jazz dance with music. Cantero et al. [51] showed an increased occipital brain activation for the higher alpha waves compared to the anterior regions. In the present study, the electrode positions O1 and O2 revealed a higher activation after the pwm and pnm conditions, which coincides with the results gained by Cantero et al. [51], except for the simultaneous decrease of the anterior brain activation that could not be confirmed. An obvious increase of the frontopolar, frontal and temporal electrode positions could be shown. These findings may rather stay in connection with the perceptions of Klimesch [52] and Hanslmayr et al. [53], who linked the alpha waves to attention and memory processes. The frontopolar and frontal electrodes (BA 10, 46, 9, 45), and therefore the frontal lobe, signify i.a. attentional processes and memory. These processes are needed for the performance of the rehearsed choreography and could be an indication for the assumption that exactly these mechanisms are activated. The electrodes T3 and T4 (BA 21, 22) may be associated with attention, memory, emotion and hearing [54, 55], which fits well as to the above presented connections. Electrodes T5 and T6 (BA 37, 39) mirror i.a. the recognition memory, which also suits the requirements of physical dance as to a rehearsed choreography with music [62]. To imagine the choreography in accordance with the music the dancer needs the recognition memory. The entire alpha frequency showed high activation across the entire scalp immediately after physically dancing, particularly, to music. Hence, it is suggested as an indication for the generation of relaxed vigilance due to physically dancing a modern jazz dance to music. This could be beneficial, especially for the creation of an optimal learning state [63].

The beta frequency band is typically related to an enhanced cortical achievement and emerges more often at the precentral and frontal brain areas [41]. It appears even in deep concentration, problem solving and fierce thinking [64, 65]. The beta waves arise not solely in mentally related processes, but also while motoric tasks, voluntary movements and permanent contractions occur [64, 65]. These perceptions may be related to the results of the current study, since the frontopolar, frontal and frontocentral electrodes achieved an increased activation after physical dancing. Most notably, these findings are true for the dancing to music, whereas the temporal and occipital brain areas additionally showed higher power in the beta frequency. The activation of the temporal electrodes, T3, T4 (BA 21/22) and T5, T6 (BA 37, 39), lasting the appearance of the beta band activity, might be a clue for conscious attention when perceiving the body in space, transforming sensory input into motoric output, recognizing patterns, feeling emotions or while hearing sounds [54, 56, 62, 66]. The occipital areas mirror specific characteristics in connection to the beta band, probably the proprioceptive attention while dancing physically and the visual appreciation of colors, shapes and movements [57]. Obviously, dancing a modern jazz dance is a motoric task and thus confirms the findings of the study by Engel and Fries [64] as well as the work of Neuper and Pfurtscheller [65], to the extent that the beta frequency and in addition the frontal lobe function are features answering to voluntary motor tasks [47]. Furthermore, it may be assumed that deep concentration immediately after physically dancing is due to the simultaneous manifestation of an increased alpha and beta power. The alpha frequency typically arises during relaxed wakefulness, which promotes deep concentration.

The occurrence of the gamma frequency is i.a. connected to movement preparation, sensomotoric and multisensoric integration [67, 68]. The modern jazz dance is composed of a complex movement combination, which requires a mellow sensomotoric and multisensoric integration to coordinate the limbs while depending on the sensory perception. Equally, movement preparation is given to initiate the rehearsed choreography. This frequency appears not only in combination with motoric tasks, but also in working memory, long-term memory and in conscious awareness [43, 69]. It could be suggested that the working memory and the conscious awareness are involved in the performance of a modern jazz dance, which can be substantial because of the empowerment of the frontal lobe (see above for the characteristics). The gamma frequency is also supposed to be involved in memory processes, if they appear in temporal brain areas [43]. This process is highly probable regarding the memorizing of the complex choreography, which increases in complexity when dancing in accordance to music is demanded.

In conclusion, the findings of this study reveal distinct brain activation across the entire scalp and for all frequency bands concerning the condition dancing physically to music and without music. Especially the alpha-2, beta and gamma frequencies were significantly higher after active dancing to music. No statistically significant time effects were analyzed for the conditions mental dancing with and without music. Differences were just shown among the physical and mental dancing condition, independent of the occurrence of music. Also, for the inter-comparison, the physical dancing conditions displayed higher activations at the frontopolar, frontal, temporal and occipital areas.

Further need for research is still required. Because dancing in western cultures is more popular with women, it would be of interest whether there are gender specific differences as to the electrical brain activity or whether there are differences in brain activity among dissimilar dance styles. To our knowledge, no studies regarding the difference of the influence of group dancing on brain activation in comparison to solo dancing exist beyond this. In this context, folk dances would be of interest for studying the impact of group dances. In sum, the topic area dance still offers lots of possibilities for research.

